# Efficient computation by molecular competition networks

**DOI:** 10.1101/2023.08.22.554117

**Authors:** Haoxiao Cai, Lei Wei, Xiaoran Zhang, Rong Qiao, Xiaowo Wang

## Abstract

Most biomolecular systems exhibit computation abilities, which are often achieved through complex networks such as signal transduction networks. Particularly, molecular competition in these networks can introduce crosstalk and serve as a hidden layer for cellular information processing. Despite the increasing evidence of competition contributing to efficient cellular computation, how this occurs and the extent of computational capacity it confers remain elusive. In this study, we introduced a mathematical model for Molecular Competition Networks (MCNs) and employed a machine learning-based optimization method to explore their computational capacity. Our findings revealed that MCNs, when compared to their non-competitive counterparts, demonstrate superior performance in both discrete decision-making and analog computation tasks. Furthermore, we examined how real biological constraints influence the computational capacity of MCNs, and highlighted the nonnegligible role of weak interactions. The study suggested the potential of MCNs as efficient computational structures in both *in vivo* and *in silico* scenarios, providing new insights into the understanding and application of cellular information processing.

## INTRODUCTION

Life senses and processes various signals to determine its response under always changing environments [1]. Compared to electronic devices, life emerges highly sophisticated computation with low energy expenditure and limited calculating elements. Researchers have focused on the high computational capacity of the human brain and found that the capacity is most endowed by the dense connections among nerve cells, which inspired the proposal of artificial neural networks (ANNs) [2]. Meanwhile, it is remarkable that even a single cell can exhibit powerful computation, such as the chemotaxis of bacteria [3] and cell-cell communication during developmental processes [4], suggesting that there are unrecognized mechanisms within cells that can enhance the efficiency of information processing.

The signal transduction network is one of the most important information processing systems within cells. In a typical signal transduction network, intracellular receptors sense extracellular ligands through binding with each other, and then the ligand-receptor complexes activate the downstream effectors. Though many ligand-receptor networks are regarded as cascade networks with specificity where one ligand binds to one receptor, ligands and receptors actually interact promiscuously in most conditions [5, 6]. The promiscuous interactions and the limited amounts arise molecular competition.

Competition can introduce crosstalk between molecules without direct physical interactions and thus forms a hidden layer of the biological network [7]. Particularly, evidences support that competition may endows efficient cellular computation. Some studies suggested that the promiscuous interactions in signal transduction networks may introduce subtle regulation [8] or enhance signal processing capacity [5, 9, 10]. For example, competition can generate diverse, non-monotonic gene regulation responses in *Escherichia coli*, where the master transcription factor cyclic AMP receptor protein (CRP) interacting with many genes simultaneously [11]. By modelling the bone morphogenetic protein (BMP) pathway quantitatively, researchers found that the promiscuous interactions in the BMP pathway serve as an effective information processing module [12]. The response pattern of the BMP pathway varies with cell-type-specific levels of receptors [12], behaving as context-dependent combinatorial logics [13] and enabling cell addressing [14].

Theoretical studies also suggested that competition may play an important role in cellular signal processing. Competition accompanied with positive feedback is able to recognize general patterns through a winner-take-all (WTA) manner [15]. By modelling a large-scale ligand-receptor system as a dynamic multiple inputs–multiple outputs system, researchers found that the system is able to efficiently sense the concentration of ligands with the temporal sequence of ligand-receptor binding and unbinding events [16]. Klumpe et al. further conceptualized the computation role of promiscuous protein interaction networks as “computational capability” and proposed a model to describe protein dimerization networks [17].

Inspired by these studies, we raised a question whether competition is the key of computation in these scenarios and how competition computes efficiently. In this study, we proposed a coarse-grained mathematical model to describe molecular competition networks (MCNs), as well as a machine learning-based parameter optimization method to efficiently explore the extent of computational capacity. We found that, in comparison with networks without competition, MCNs as well as some kinds of their variants can well perform both discrete decision-making and analog computation tasks. We also modeled kinds of constraints in real biological scenarios and revealed how these constraints affect the computational capacity of MCNs. We found that weak interactions in MCNs play a nonnegligible role in computation and quantitatively revealed that a high sensitivity of dose response can be achieved by competition. We further proposed that MCNs may serve as efficient computation structures in both synthetic circuits and pattern recognition tasks. In summary, this study highlights the critical role of competition in cellular computation and offers valuable insights and methodologies for harnessing MCNs both *in vivo* and *in silico*.

## METHODS

### The coarse-grained model of MCNs

Almost all molecules in cells participate in more than one biochemical reaction. The amount of these molecules can hardly be regarded as unlimited. Therefore, competition for these molecules arises between different reactions. Here we considered a competition system with two classes of molecules (Fig. 1a), noted as *A*_*i*_ (*i* = 1, …, *n*_*A*_) and *B*_*j*_ (*j* = 1, …, *n*_*B*_), respectively. Different from specific binding, each *A*_*i*_ may reversibly bind with each *B*_*j*_ to form *C*_*ij*_:

**FIG. 1.**
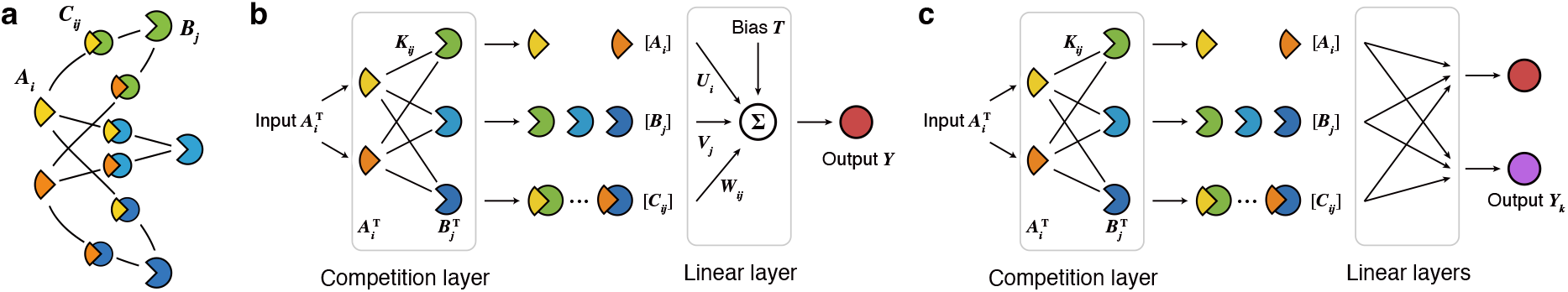
(a) The schematic diagram of molecular competition. (b) The illustration of MCNs with one output. (c) The illustration of MCNs with multiple outputs.

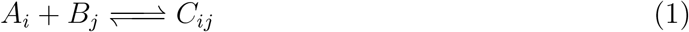

We assumed the binding reaction usually happens at a smaller time scale compared to other cellular processes. We ignored the production and degradation of these molecules and only focused on the steady state of the system in this study. We used 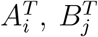 and 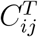 to denote the total concentration of *A*_*i*_, *B*_*j*_ and *C*_*ij*_, and used [*A*_*i*_], [*B*_*j*_] and [*C*_*ij*_] to denote the equilibrium concentration of *A*_*i*_, *B*_*j*_ and *C*_*ij*_.

When every reaction comes to equilibrium, the system shows

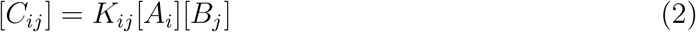

where *K*_*ij*_ is the equilibrium constant of Reaction 1.

By the conservation of mass, we got

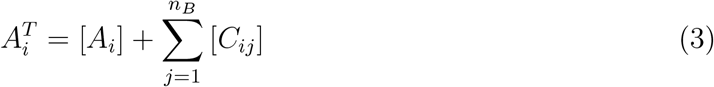

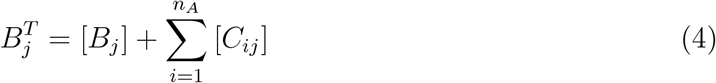

When *n*_*A*_ *>* 1 and *n*_*B*_ = 1, the system degenerates into a multi-to-one competition, which was well characterized in our previous study [7].

The equilibrium concentration of each substance is determined by the total concentration of each substance and the equilibrium constants. From the perspective of information processing, if we take all 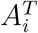 as inputs, all [*C*_*ij*_] as outputs, and all *K*_*ij*_ and 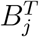 as parameters, the competition system performs like a computation module. Similar to layers in ANNs, We named it as a “competition layer” (Fig. 1b):

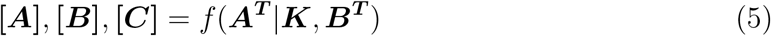

Referring to biological signaling transduction networks, we supposed each substance has a linear effect on downstream signal *Y* with weights *U*_*i*_, *V*_*j*_, *W*_*ij*_ and a bias *T* . It is the same as the “linear layer” in ANNs (Fig. 1b):

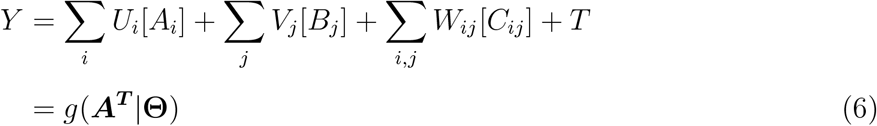

where **Θ** denotes the parameter sets of the model {***K, B***^***T***^, ***U***, ***V***, ***W***, *T*}. A competition layer and a linear layer compose the coarse-grained model of MCNs (Fig. 1b). By changing the linear layer to multiple outputs *Y*_*k*_ (*k* = 1, …, *n*_*Y*_), MCNs can easily deal with more complex tasks such as classification (Fig. 1c).

### Solving the steady state of the competition layer

According to the Deficiency Zero Theorem [18], if a chemical reaction network is deficiency zero and weakly reversible, and obeys mass action kinetics, there always exists precisely one equilibrium which is asymptotically stable. Thus, there is and only one steady state of the competition layer. As solving the steady state of a dynamic system by numerical simulation is usually time-consuming and complicated, here we proposed a simple iterative method to calculate the steady state of the competition layer. By joining Eqs. 2, 3 and 4, we got

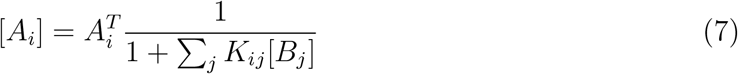

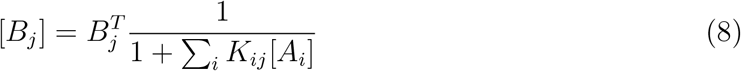

which describes the partition of molecules between free and bound states. We initialized all [*A*_*i*_] with 0, then alternately calculated [*B*_*j*_] and [*A*_*i*_] by Eqs. 7 and 8 until converging. This process can also be regarded as a simplification of the concentration allocation between different forms of molecules in discrete time steps.

### Parameter constraints

Parameters of biological models always obey strict constraints to be in line with reality, where most realistic parameters are non-negative. For example, the total concentration 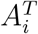 and 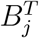 and the equilibrium constant *K*_*ij*_ must be non-negative to have a physical significance. In some scenarios which we discussed more in Results, the weights *U*_*i*_, *V*_*j*_, *W*_*ij*_ and the bias *T* of the linear layer should also be non-negative. We formulated each non-negative parameter as *x* = exp(*x*^*∗*^) and regarded *x*^*∗*^ as a model parameter to be optimized. This reparametrization trick also allowed us to explore a larger parameter space, as biological parameters usually have a wide range of variation.

### Machine-learning-based parameter optimization

Functions of a biological network highly rely on its parameter settings. A lot of studies tried to explore a mass of parameters through scanning [7] or random sampling [19] to describe the function of biological networks, but these approaches are not efficient when the parameter space is high-dimensional. We transformed this exploration-based approach into a learning-based approach, i.e. first giving all the functional targets *g* in Eq. 6 that we care about, and then optimizing **Θ** to evaluate whether the goals can be achieved.

Machine learning algorithms have been implemented to optimize the topology or parameters of biological networks in previous studies [20, 21]. Here we adopted a plain machine-learning-based approach to optimize the parameters of MCNs. We performed the gradient descent algorithm to reduce the difference between the model output and the target to find a local minimum of a loss function. The gradients were calculated by the automatic differentiation technique. As the model is likely to converge to a local minimum, we repeated the training process ten times with different initialization settings and chose the model with the best performance for further analysis.

We implement all the above-mentioned approaches in Python. The code is available at https://github.com/maplecai/competitve_network.

## RESULTS

### The computational capacity of unconstrained MCNs

Discrete decision-making such as Boolean operations are the fundamental functions in general information processing, DNA computing, and classical synthetic gene circuits [15, 22, 23]. Thus, we chose 2-input Boolean functions as the target functions to evaluate the computational capacity of MCNs. We also selected 3-input Boolean functions and 2-input 3-quantized Boolean functions as tasks to investigate whether MCNs could process multiple inputs or non-digital inputs (Fig. 2a). For 2-input Boolean functions, we considered the input 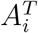 at 10^*−*1^ (low) or 10^1^ (high), forming 2^2^ = 4 input tuples. Each output could be either 0 (low) or 1 (high), so there are 2^4^ = 16 possible functions in total. Similarly, for 3-input Boolean functions, we set 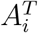 at 10^*−*1^ (low) or 10^1^ (high), forming 2^3^ = 8 input tuples and 2^8^ = 256 different functions. As for 2-input 3-quantized Boolean functions, we set 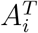 at 10^*−*1^ (low), 1 (medium), or 10^1^ (high), forming 3^2^ = 9 input tuples and 2^9^ = 512 different functions.

**FIG. 2.**
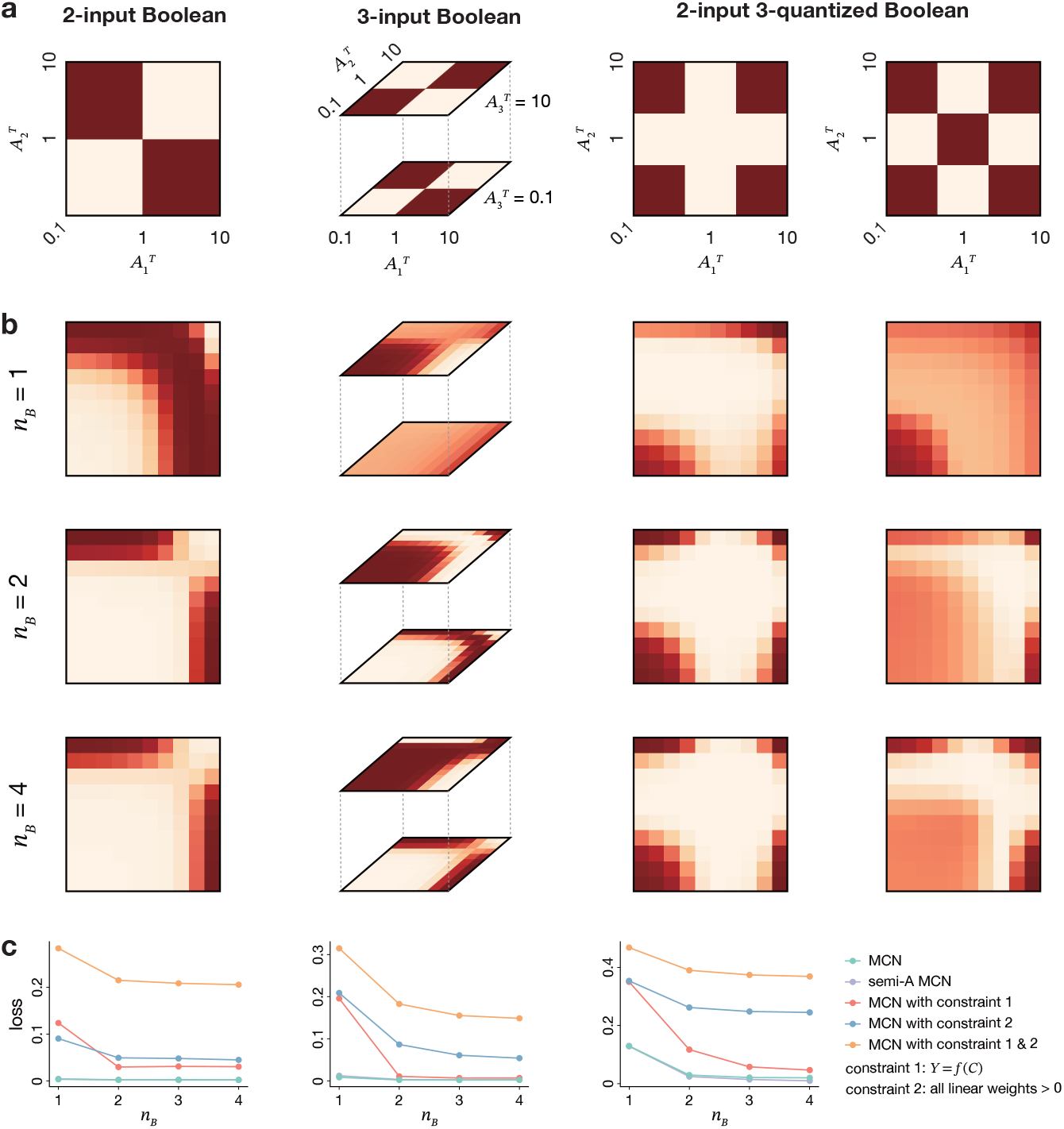
The schematic diagram of target functions and model outputs. (a) Visualization of some target functions in different tasks. (b) Visualization of model outputs that fit the targets functions fitted by MCNs with *n*_*B*_ = 1, 2, 4. (c) The average cross-entropy loss of different models.

We optimized the parameters of a MCN to make it best performs a desired Boolean operation. Since the outputs are discrete, we treated the goal as a classification task. Parameters were optimized to minimize the cross-entropy loss to evaluate the difference between the model output and the target output. After training, if there was a threshold according to which the model output could be quantized to be the same as the target function, we regarded the training result as a successful fit. Besides, we visualized the model outputs on a 10*×*10 grid within the range of inputs at the logarithmic scale to investigate its computation pattern (Fig. 2b).

We selected a linear model 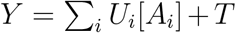 as the baseline model. When there doesn’t exist any competition effect in a MCN, Eqs. 3 and 4 degenerates into 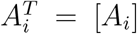 and 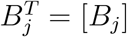 and thus Eq. 6 is equivalent to the linear model. The linear model describes the computation directly driven by inputs 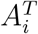 without any intermediate information processing.

This model is familiar in gene regulation modelling to describe a gene regulated by multiple factors without any antagonistic or synergistic effect. There are 14 (88%), 104 (41%) and 60 (12%) linear separable functions in three different tasks, respectively. As shown in Table I, the linear model perfectly fits every linear separable functions, but cannot fit any linear inseparable functions. For instance, in the 2-input Boolean function task, XOR and XNOR are the two functions that cannot be fitted, which are both linear inseparable functions.

**TABLE I.**
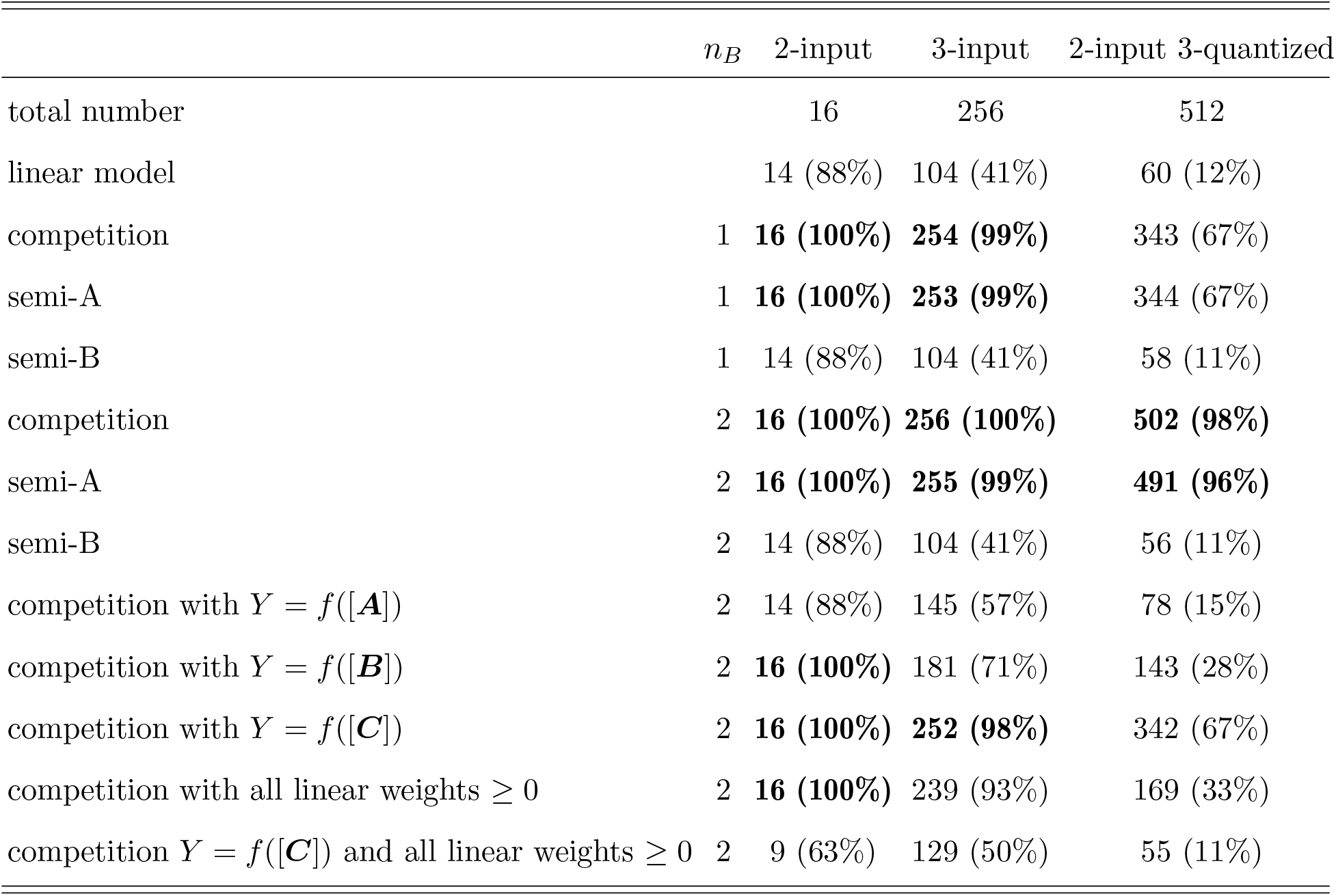
Number of successfully fitted functions.

By contrast, we found that MCNs show great performance on these tasks (Table I). MCN, even with only one type of *B* (*n*_*B*_ = 1), can perfectly fit all the 2-input Boolean functions, even for the linear inseparable functions XOR and XNOR (Fig. 2b). The good performance may explain the role of co-receptors in many signaling pathways [24–26], which can be regarded as an competition layer with *n*_*B*_ = 1.

MCNs can also performs well in the 3-input Boolean function tasks. MCNs with *n*_*B*_ = 1 can fit 254 (99%) functions, and MCNs with *n*_*B*_ = 2 can fit all 256 (100%) functions. The two failed functions with *n*_*B*_ = 1 are 3-input XOR and XNOR. We retrained the model for more times on these two functions, but MCNs with *n*_*B*_ = 1 consistently failed to fit them, suggesting that these tasks exceed the computational capacity of MCNs with *n*_*B*_ = 1 but can be achieved with *n*_*B*_ = 2 (Fig. 2b).

The 2-input 3-quantized Boolean function task is more challenging. MCNs with *n*_*B*_ = 1 can fit 343 (67%) functions, and MCNs with *n*_*B*_ = 2 can fit 502 (98%) functions. Some of the failed functions can be achieved when *n*_*B*_ increases. But there are still two patterns that can’t be fitted when *n*_*B*_ increases to 4. One pattern is showed in the right panel of Fig. 2a, and the other pattern is its inversion. We supposed that the complexity of these two patterns exceed the computational capacity of MCNs with any *n*_*B*_.

We also investigated the cross-entropy loss of these tasks. We found that the loss of MCNs is significantly lower than that of the linear model. The loss of MCNs gradually decreases with the increase of *n*_*B*_ in all tasks and tends to be saturated when *n*_*B*_ *≥* 2 for 2-input Boolean and 3-input Boolean functions, and *n*_*B*_ *≥* 4 for 2-input 3-quantized Boolean functions (Fig. 2c). All the results suggested the high computational capacity of MCNs in dealing with discrete decision-making tasks, which is particularly evident when facing linear inseparable functions.

### MCNs with some variants keep efficient computation

Particular molecular competition scenarios may exhibit some differences. Our MCN model can describe various competition-related systems with minor changes. In some scenarios, the binding of one molecule to a target may not inhibit the binding of another molecule to the target. For instance, in the microRNA(miRNA)-mRNA competition system [27, 28], different miRNAs can bind with a mRNA at different sites, and thus the bindings of these miRNAs are not mutually exclusive. To model this system, we set the concentration of molecule *A* or *B* as a fixed value. For example, when considering *A* as fixed, we replaced Eq. 3 into 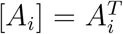and named this system as a semi-A MCN. Similarly, we defined a semi-B MCN by changing Eq. 4 to 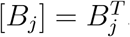 . However, in a semi-B MCN, Eq. 3 degenerates into

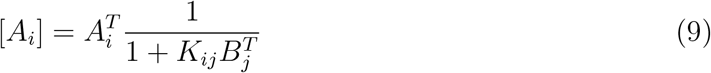

and the model is equivalent to the linear model.

We investigated the performance of semi-A and semi-B MCNs with the same methods (Table I). As expected, semi-B MCNs perform the same as the linear model. Interestingly, semi-A MCNs perform well in all tasks, just slightly inferior than MCNs in 3-input and 2-input 3-quantized Boolean tasks. The cross-entropy loss of these models supports the above conclusions (Fig. 2c). The results suggested that the competition of *B* is the key of nonlinear computation in MCNs when considering *A*^*T*^ as inputs.

### MCNs with biological constraints

Any MCN in real world should obey biological constraints. Understanding the behavior of MCNs with biological constraints is necessary to reveal their biological roles and construct synthetic competition networks with desired functions. The above-mentioned model imposed only the necessary constraint that parameters in the competition layer must be non-negative. Here we considered two optional constraints: (1) only one species of molecules (**[*A*], [*B*]** or **[*C*]**) can drive the downstream signal *Y* ; (2) all weights in the linear layer must be non-negative. Both the constraints are familiar in many biological systems. For example, in ligand-receptor systems, the active molecules are sometimes the ligand-receptor complexes *C*_*ij*_, sometimes the receptors *B*_*j*_, but rarely *A*_*i*_, *B*_*j*_, and *C*_*ij*_ at the same time. Furthermore, the downstream effects of these molecules are usually similar, thus leading to the same sign of their weights in the linear layer.

For the first constraint, we found that the constraints of *Y* = *f* ([***A***]) and *Y* = *f* ([***B***]) significantly affect the performance of the model. The model with *Y* = *f* ([***A***]) constraints could only fit about 57% 3-input and 15% 2-input 3-quantized Boolean functions, and 71% and 28% when *Y* = *f* ([***B***]) constraints (Table I). However, the constraint of *Y* = *f* ([***C***]) showed less effects on computational capacities of MCNs. The performance decreased very little from 100% to 98% in 3-input functions, and from 98% to 67% in 2-input 3-quantized functions (Table I). The results showed that the constraint of *Y* = *f* ([***C***]) is much weaker than the constraints of *Y* = *f* ([***A***]) or *Y* = *f* ([***B***]).

The second constraint affect the performance little. We found that the model could still perfectly fit 100% the 2-input Boolean functions, but the ability of fitting 3-input and 2-input 3-quantized functions is much lower (93% and 33%, respectively). Adding *n*_*B*_ are somehow useless. It could fit 254 (99%) 3-input functions, but only 187 (37%) 2-input 3-quantized functions when *n*_*B*_ = 4.

When both constraints exist, the performance of MCNs decreases significantly in all tasks. The observations were similar when considering the cross-entropy loss (Fig. 2c). It should be noted that in this condition, when all inputs are set as low, the output will definitely exhibit low. Therefore, the model can theoretically fit up to about half of functions in any task. In fact, the model can fit 129 3-input functions, containing half of the total as well as an always-TRUE function, which is its theoretical limit. If we were allowed to reverse input signals before the model, MCNs could fit up to 246 (96%) functions. All the results suggested that MCNs could remain strong computation capability even when constraints are applied in real biological scenarios.

### Roles of weak interactions

We found that the model with both constraints could fit the XOR function, similar to the “imbalance” pattern observed by Antebi et al [12]. Here we illustrated the parameter settings of the MCN that generates the XOR function, with *n*_*B*_ = 2, *Y* = *f* ([***C***]), and all linear weights *≥* 0 (Fig. 3a). We found that *C*_12_ and *C*_21_ showed low binding affinity but high output efficiency. On the contrary, *C*_11_ and *C*_22_ showed high binding affinity but low output efficiency. It supports the previous results about anti-correlations between the affinity and activity found by Su et al [14].

**FIG. 3.**
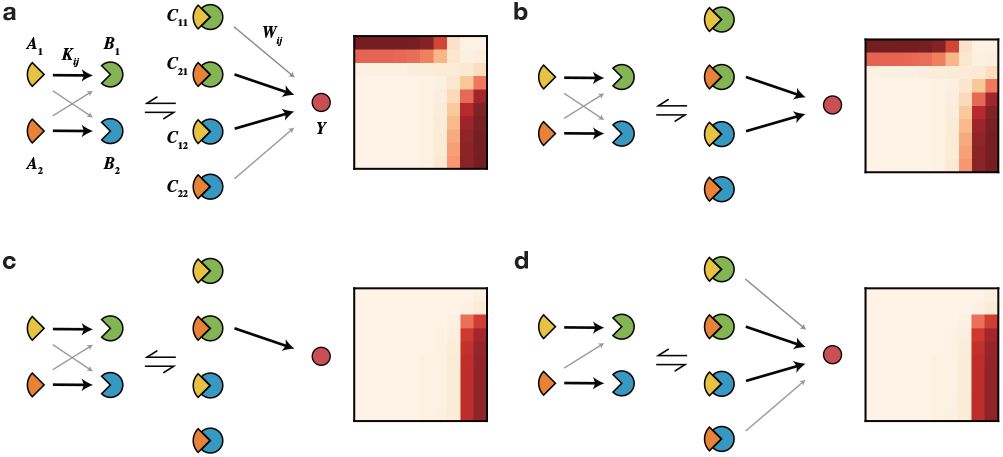
Parameter settings and model outputs of MCNs that fit the XOR function. For all MCNs, *n*_*B*_ = 2, *Y* = *f* ([***C***]), and all linear weights *≥* 0. The width of the line represents the strength of the interaction. Results of MCNs with (a) no modification; (b) two weakest linear activity parameters set as zero; (c) three weakest linear activity parameters set as zero; (d) the smallest competition affinity parameter set as zero.

We wondered whether weak relationships in MCNs are necessary. We removed interactions with low weights and investigated how output *Y* changes. When we set two weak linear activity parameters *W*_*ij*_ as zero, the output of the model was almost unchanged (Fig. 3b). When setting an additional *W*_*ij*_ as zero, the output failed to generate the XOR function (Fig. 3c). The results suggested that multiple components in the linear layer is essential for generating complex patterns, but it is not a must to be dense.

Interestingly, when we set the smallest competition affinity parameter *K*_*ij*_ as zero, the model failed to generate the XOR function (Fig. 3d). In this condition, the system cannot generate any *C*_*ij*_ and thus is similar to setting the corresponding *W*_*ij*_ as zero. Actually, in most training results, the activity *W*_11_ and *W*_22_ in our model is exactly zero, but the affinity *K*_12_ and *K*_21_ is strictly greater than zero. All the results suggested that the affinities *K*_*ij*_, even weak affinities, play important roles in computation. We further investigated the number of non-zero linear activity parameters *W*_*ij*_ in MCNs of other target functions, and found that at most two active molecules *C* is enough for any 2-input Boolean function. It indicates that the linear layer in MCNs is always sparse, which is helpful for biological implementation.

### Analog computation with MCNs

In the above-mentioned cases, we only considered the output under binarized or ternarized inputs. However, biological systems usually adopt analog information processing as the concentrations of molecules always change continuously [29, 30]. Thus, we tried to investigate whether MCNs can well fit analog-input analog-output functions.

This task can be regarded as a regression task. We considered the 2-input Boolean functions. We took output 0 (low) or 1 (high) as real values instead of class labels and generated 10 *×* 10 grid as the input at the logarithmic scale. We then interpolated the output to the grid input using two approaches: the nearest neighbor interpolation algorithm to achieve a Boolean-like target function, and the bilinear interpolation algorithm to achieve a continuous target function (Fig. 4a). We trained MCNs without constraints and adopted the mean squared error (MSE) as the training loss.

**FIG. 4.**
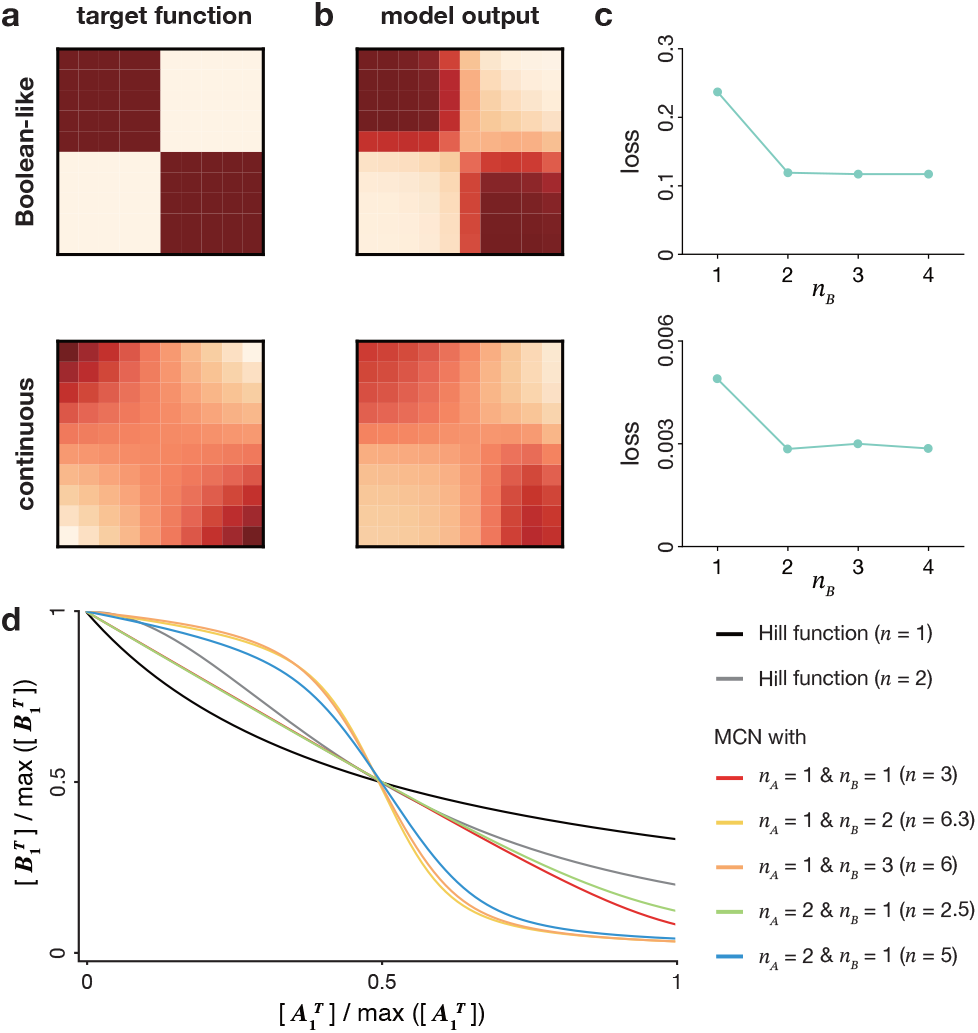
MCNs trained by 10 *×* 10 grid inputs. (a-b) Visualization of the Boolean-like target function (a) and the continuous target function (b) of XOR and the fitted results of MCNs with *n*_*B*_ = 2. (c) The mean MSEs of MCNs with different *n*_*B*_ when fitting Boolean-like target functions and continuous target functions. (d) The dose-response curve of different kind of MCNs fitting the step function. *n* denotes the fitted Hill coefficient.

As shown in Fig. 4b, MCNs with *n*_*B*_ = 2 could well generate XOR functions, which is much similar to the target XOR function. In general, MCNs with *n*_*B*_ = 2 could well fit all 16 patterns of Boolean-like target functions with a mean MSE *<* 0.03, and all patterns of continuous target functions with a mean MSE *<* 0.006 (4b). The mean MSE didn’t decrease with the increase of *n*_*B*_. These results showed that MCNs can well fit analog functions, and the training approaches are compatible to analog targets without any modification.

### Competition sharpens dose-response curve

As shown in Fig. 4b, MCNs can generate shape edges when fitting Boolean-like functions. It is consistent with previous studies on competition systems with *n*_*A*_ = 1 that competition can perform a very sharp transition in dose-response curve to achieve ultrasensitivity [7, 31– 33]. However, we didn’t know the upper bound of ultrasensitivity that MCNs can achieve and whether more species of *A* could generate higher sharpness. Here we take 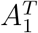 as the input and 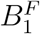 as the output, with constraints of *Y* = *f* ([***C***]) and linear weights *≥* 0. We set the target function as a step function with input range between 0 and 2 and regarded it as an analog function. We trained MCNs to fit the function, and then calculated the Hill coefficient of the model’s output as the sharpness metric [34].

The fitted Hill coefficients of semi-A MCNs are always equal to 1. As shown in Fig. 4d, the coefficient of MCNs is larger than 1, indicating that that competition can indeed generate a sharper curve. When *n*_*B*_ = 2, the Hill coefficients significantly increase up to around 6 but no longer grow with the increase of *n*_*B*_. We also found that the Hill coefficient is only determined by *n*_*B*_ instead of *n*_*A*_. These results showed the upper bound of the ultrasensitivity that a MCN can achieve, and the machine learning-based method can help to optimize the design of parameters when desiring a high sharpness of the dose-response curve.

### MCNs in pattern recognition tasks

The above results suggested that MCNs may play important roles in cellular functions. We further explored the performance of MCNs without any constraints in acting multi-class classification tasks. We first applied MCNs on the Iris dataset [35], which is a classic machine learning dataset. It contains a total of 150 samples in 3 classes, and each sample has 4 features. As mentioned before, MCNs could have multiple outputs by increasing *n*_*Y*_ (Fig. 1c). Thus, we established MCNs with *n*_*A*_ = 4 and *n*_*Y*_ = 3. Each *A* corresponds to a feature and each *Y* corresponds to a class. The training objective is to maximize the output *Y* node of the correct class and minimize the others. We set *n*_*B*_ from 1 to 4 to test the performance of MCNs with different *n*_*B*_. The MCNs got accuracies around 97% regardless of *n*_*B*_, which is close to the 97% accuracy of logistic regression.

We also trained MCNs on the MNIST dataset [36] which is much bigger than Iris. MNIST dataset contains 60,000 handwritten digit images of 10 hand-written digits, and each image has 28 *×* 28 pixels and 1 channel. It is a 784-input 10-output problem, so we set MCNs with *n*_*A*_ = 784 and *n*_*Y*_ = 10 to handle it. MCN could achieve 80% accuracy when *n*_*B*_ = 1 and 87% accuracy when *n*_*B*_ *>* 2. As a comparison, logistic regression has 93% accuracy. The results demonstrated that MCNs could also perform *in silico* pattern recognition tasks.

## DISCUSSION

Competition is inevitable in biological systems. In this study, we showed that competition introduces nonlinear interactions and thus benefits computational capacity of biological networks. These results provided clues to reveal how cells process complicated signals with limited genes and resources with inherent competition. We inferred that competition may be a foundation of information processing within cells rather than a frustrated phenomenon to be avoided. There are several aspects to be investigated in future to further illustrate the role of MCNs in the competition capacity of cells, such as dynamic signaling processing and encoding, energy consuming, robustness, and cell-type-specific function switching.

We supposed that MCNs could be abstracted as a general network structure for *in silico* applications. Actually, the structure of MCNs is a bipartite graph, quite similar to the restricted Boltzmann machine (RBM) [37]. RBM is a classic ANN that considers the probability distribution of each node according to its “energy”. Similarly, MCNs are natural systems that depict the probability distribution of each molecule according to the energy of all chemical reactions. Thus, we believe that MCNs may be close to or even better than RBM in machine learning.

It should be noticed that MCNs are not the same as the WTA structure that exists in many biological processes [38, 39] and has been widely used in the design of ANNs [40, 41], DNA computing [15, 22, 23, 42, 43], and synthetic gene circuits [44]. WTA behavior raises when competition is accompanied with positive feedback loops [15]. Actually, WTA can be regarded as a simplification of MCNs with only one *B*_*j*_≠ 0. This simplification leads to a digital-like computation manner, which makes the system easy to be analyzed but could cause extra consumption of energy and molecules [30, 45]. The MCN model is a closer model to real biological networks with richer properties and can be easily extended to analyze various behaviors including WTA.

MCNs have the potential to be applied in constructing synthetic gene circuits to simplify the structures. Traditional synthetic Boolean operation modules in synthetic gene circuits are usually assembled by some basic logic operation units, which was inspired by electronic circuits. However, such approach would lead to high complexity and difficult implementation. For instance, when using standardized NOT and NOR gates to build Boolean operation modules, a 2-input XNOR module should contain 4 gates, and a 2-input Consensus module (outputting high when three inputs are the same) should contain seven gates [46]. MCNs can accomplish both tasks with *n*_*B*_ = 1. Actually, MCNs can perform all 2-input Boolean functions with *n*_*B*_ = 1 and all 3-input Boolean functions with *n*_*B*_ *≤* 2. This may greatly simplify the design approach and save the energy and resource consumed by the synthetic gene circuits. With the rapid development of AI-aided design of DNA, RNA, and protein, MCNs may be easily implemented in synthetic biology and thus revolutionize future synthetic gene networks.

In this work, we proposed a machine-learning-based method to optimize parameters in biological networks. Different from the exploration-based approach, this approach provides an opposite way for studying complex biological networks: first hypothesizing what function the networks can achieve, and then setting the function as a goal for parameter optimization. We found that the optimization approach proposed in this work is not robust and easily falls into a local minimum. More should be investigated to obtain a stable strategy for optimizing highly nonlinear biological networks.

The work was supported by the National Natural Science Foundation of China (grants 62250007, 62225307, 62103227, and 61721003), and the State Key Research Development Program of China (grant 2020YFA0906900).

H.C. and L.W. contributed equally to this work.

